## Supplementary Material for "Steering clear of the confusion effect: aerial predators target fixed points in dense prey aggregations"

This PDF contains:

Figures S1-S3
Tables S1-S3
Legend for Movie S1
References

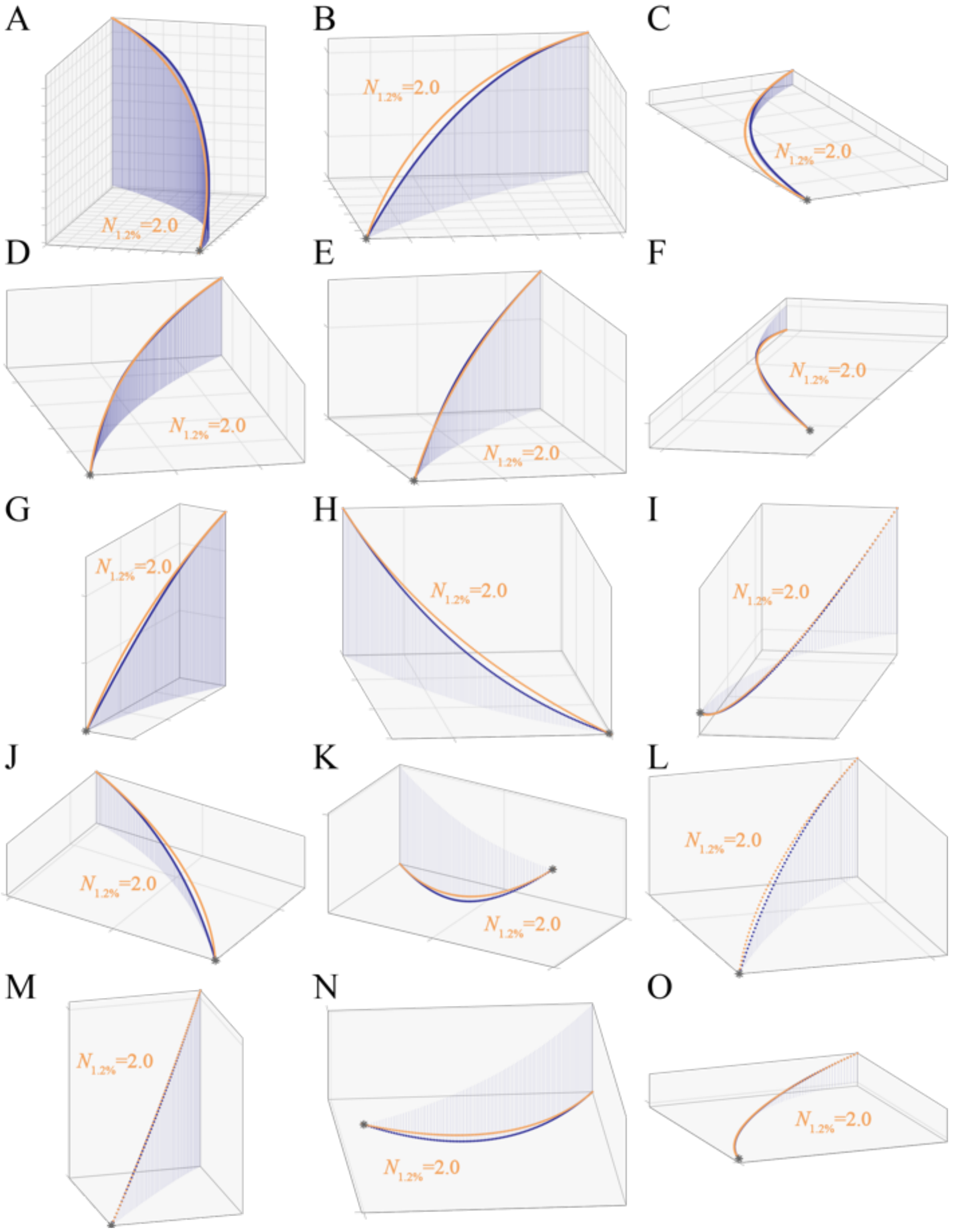


**Figure S1. Long-range approaches of Swainson's Hawks attacking Mexican Free-tailed Bats modelled as constant radius turns into the swarm. (A-O)** Each panel plots the reconstructed three-dimensional attack trajectory of an incoming hawk (dark blue points); dark blue lines are dropped vertically from each point to accurately convey the three-dimensional shape of the trajectory. It was not possible to track the bats that the hawks grabbed at this range, but orange lines plot simulations of the hawk's flight trajectory generated under delay-free PN guidance at a fixed navigation constant of $N = 2$, assuming flight at the same speed as measured and treating the final position of the hawk as the target. This serves to generate a constant radius turn that satisfies the kinematic constraint of passing through the hawk's initial and final positions whilst also matching its initial flight velocity. Trajectories are plotted for the longest section of flight for which the relative error remains below the threshold value of $\varepsilon \leq 0.012$, displaying the subset of 15 flights with the longest simulations meeting this criterion. Grid spacing: 10 m.

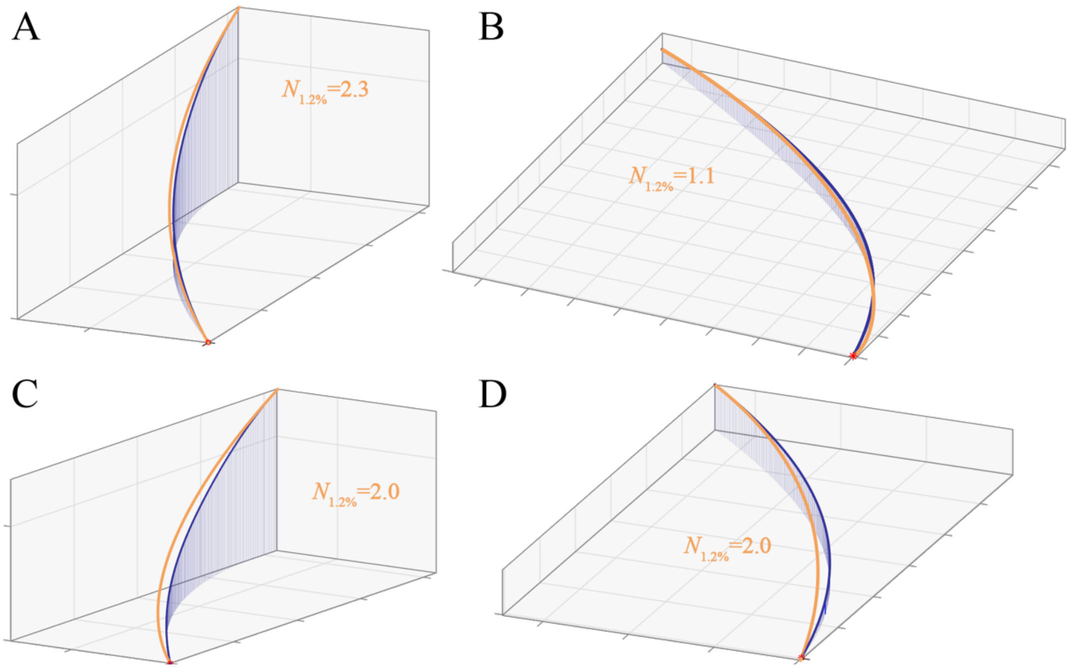


**Figure S2: Long-range approaches of Peregrine or Prairie Falcons attacking Mexican Free-tailed Bats modelled under delay-free PN targeting a fixed point in the swarm.** Each panel plots the reconstructed three-dimensional attack trajectory of an incoming falcon (dark blue points), where panels (A,C) and (B,D) represent the same trajectory fitted under a different guidance model. Dark blue lines are dropped vertically from each point to accurately convey the three-dimensional shape of the trajectory. It was not possible to track the bats that the falcons grabbed at this range, but orange lines plot simulations of the falcon's flight trajectory generated under delay-free PN guidance at: **(A,B)** the best-fitting value of the navigation constant $N$; **(C,D)** a fixed navigation constant of $N = 2$. The latter serves to generate a constant radius turn that satisfies the kinematic constraint of passing through the hawk's initial and final positions whilst also matching its initial flight velocity. These simulations assume flight at the same speed as measured and treat the falcon's final position as the target. Trajectories are plotted for the longest section of each flight for which the relative error remains below the threshold value of $\varepsilon \leq 0.012$. Grid spacing: 10 m.

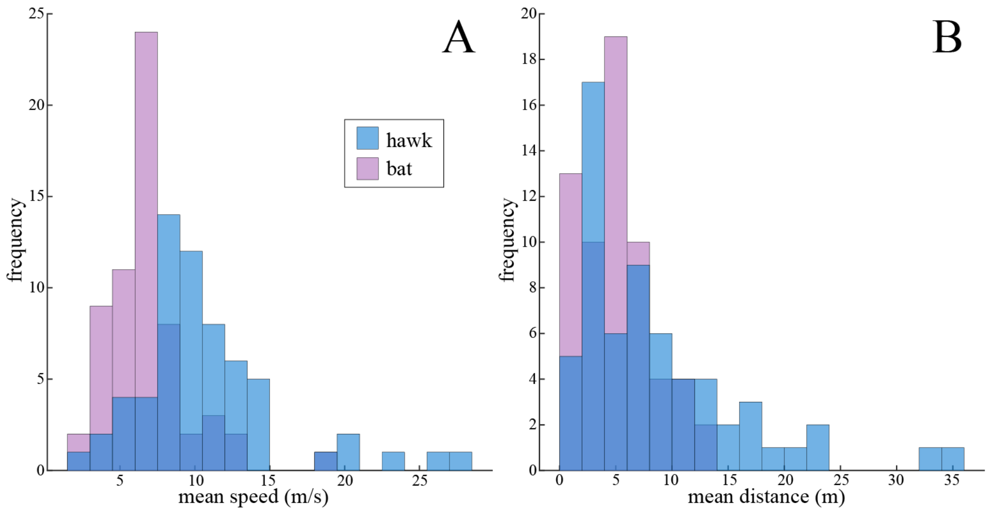


**Figure S3. Histograms summarizing measured flight performance of hawks and bats by flight. (A)** Mean flight speed. **(B)** Mean path length. Histograms display data for all $n = 62$ terminal attack trajectories.

**Table S1. Summary of fitted guidance models.** Performance of delay-free guidance models fitted to the $n = 62$ terminal attack trajectories of Swainson's Hawks attacking swarming Mexican Free-tailed Bats, for the different combinations of guidance law (i.e. PN, PP or PN+PP) and target definition (i.e. instantaneous bat position, final bat position, final hawk position). Tilde notation denotes the median values of the guidance constants $N$ or $K$, and relative error $\varepsilon$, together with the corresponding bias-corrected and accelerated bootstrap 95% confidence interval (CI).

| **target of guidance** | **fitted guidance law** | | | | | | |
|---|---|---|---|---|---|---|---|
| | **PN** | | **PP** | | **PN+PP** | | |
| | $\tilde{N}$ **(95% CI)** | $\tilde{\varepsilon}$ **(%) (95% CI)** | $\tilde{K}$ **($s^{-1}$) (95% CI)** | $\tilde{\varepsilon}$ **(%) (95% CI)** | $\tilde{N}$ **(95% CI)** | $\tilde{K}$ **($s^{-1}$) (95% CI)** | $\tilde{\varepsilon}$ **(%) (95% CI)** |
| **instantaneous bat position** | 1.63 (1.08, 2.04) | 2.08 (1.67, 2.68) | 0.66 (0.13, 1.62) | 2.97 (2.34, 4.15) | 1.83 (1.17, 2.57) | 0.36 (-0.29, 0.66) | 0.88 (0.64, 1.09) |
| **final bat position** | 1.65 (1.45, 2.07) | 1.09 (0.89, 1.34) | 3.06 (2.71, 4.03) | 1.23 (1.03, 1.57) | 1.96 (1.28, 3.16) | 0.24 (-1.10, 0.54) | 1.04 (0.80, 1.22) |
| **final hawk position** | 1.91 (1.81, 2.01) | 0.04 (0.03, 0.08) | 4.36 (3.51, 5.13) | 0.30 (0.24, 0.43) | 2.08 (2.03, 2.16) | -0.77 (-1.17, -0.41) | 0.01 (0.00, 0.01) |

**Table S2. Validation of the principle of using model selection to identify the target of PN guidance.** Re-analysis of published data for Peregrines and Gyrfalcons pursuing singleton aerial targets [1, 2], showing the results of fitting delay-free PN guidance targeting either the instantaneous or final position of the target. To quantify how well each model fits the data, we identify the path length and duration of the longest section of each flight for which the relative error remains below the threshold value of $\varepsilon \leq 0.012$, up to and including the point of intercept. As expected, the data are better modelled by taking account of the full information on target position, in contrast to our results for Swainson's Hawks attacking swarming bats.

| **target of PN guidance** | **median path length modelled (m) (Q1, Q3)** | | **median duration modelled (s) (Q1, Q3)** | |
|---|---|---|---|---|
| | **Peregrines** | **Gyrfalcons** | **Peregrines** | **Gyrfalcons** |
| **instantaneous target position** | 105.8 (44.8, 194.3) | 35.1 (21.6, 89.6) | 5.4 (3.1, 11.6) | 4.6 (3.2, 8.2) |
| **final target position** | 86.4 (53.4, 104.9) | 27.1 (21.0, 36.1) | 4.7 (4.1, 5.5) | 3.6 (2.8, 5.2) |

**Table S3. Contingency table summarizing success of attacks by Swainson's Hawks on swarming Mexican Free-tailed Bats.** Bats classified as flying outside the swarm column were judged to be flying > 5 body lengths from their nearest neighbour and/or appeared to be flying in a different direction to its coordinated members. Bats not meeting these criteria were classified as flying within the swarm column. Attacks are classified as successful if the hawk succeeded in grabbing the bat in its talons.

| | successful captures | failed captures | *totals* |
|---|---|---|---|
| attacks on bats flying outside swarm column | **3** | **5** | *8* |
| attacks on bats flying within swarm column | **12** | **42** | *54* |
| *totals* | *15* | *47* | *62* |

**Movie S1.** Sample video footage of two Swainson's Hawks *Buteo swainsoni* attacking swarming Mexican Free-tailed Bats *Tadarida brasiliensis*. Video recorded in full HD (1920×1080 pixels) at 50 Hz frame rate. The three-dimensional trajectories of the hawks and bats were reconstructed by tracking them across stereo video pairs (see Methods).
